# Structure of the poxvirus decapping enzyme D9 reveals its mechanism of cap recognition and catalysis

**DOI:** 10.1101/2021.10.04.463123

**Authors:** Jessica K. Peters, Ryan W. Tibble, Marcin Warminski, Jacek Jemielity, John D. Gross

**Affiliations:** Department of Pharmaceutical Chemistry, University of California, San Francisco, San Francisco, CA, USA; Program in Chemistry and Chemical Biology, University of California, San Francisco, San Francisco, CA, USA; Division of Biophysics, Institute of Experimental Physics, Faculty of Physics, University of Warsaw, Warsaw, Poland; Centre of New Technologies, University of Warsaw, Warsaw, Poland

## Abstract

Poxviruses encode decapping enzymes that remove the protective 5’ cap from both host and viral mRNAs to commit transcripts for decay by the cellular exonuclease Xrn1. Decapping by these enzymes is critical for poxvirus pathogenicity by means of simultaneously suppressing host protein synthesis and limiting the accumulation of viral dsRNA, a trigger for antiviral responses. Here we present the first high resolution structural view of the vaccinia virus decapping enzyme D9. This Nudix enzyme contains a novel domain organization in which a three-helix bundle is inserted into the catalytic Nudix domain. The 5’ mRNA cap is positioned in a bipartite active site at the interface of the two domains. Specificity for the methylated guanosine cap is achieved by stacking between conserved aromatic residues in a manner similar to that observed in canonical cap binding proteins VP39, eIF4E, and CBP20 and distinct from eukaryotic decapping enzyme Dcp2.

## INTRODUCTION

Poxviruses are large, double-stranded DNA viruses that replicate and assemble infectious particles exclusively in the cytoplasm of cells (Moss, 2001). Although poxviruses encode approximately 200 proteins that are expressed in sequential stages of viral replication, they rely on cellular translation machinery for viral protein synthesis (Moss, 2001). To mitigate competition with host mRNAs and simultaneously reduce the synthesis of innate and adaptive immune response factors, poxviruses induce a rapid decline in host protein synthesis through several mechanisms, including mRNA decay. Like eukaryotic mRNAs, poxvirus mRNAs contain a 5’-7-methylguanosine (m^7^G) cap and 3’-poly(A) tail. In contrast to host mRNAs, poxvirus transcripts are modified in the cytoplasm by viral enzymes (Broyles and Moss, 1987). The 5’ cap protects the mRNA from exoribonuclease digestion as well as allows for cap-dependent translation of viral mRNAs (Cantu et al., 2020). Although seemingly counterintuitive, poxviruses also encode decapping enzymes that remove the protective m^7^G cap producing m^7^GDP and a 5’ monophosphate RNA which is then committed to decay by cellular 5’-3’ exonuclease Xrn1 (Burgess and Mohr, 2015; Parrish and Moss, 2007; Parrish et al., 2007).

Vaccinia Virus (VACV), the prototypic poxvirus, encodes two decapping enzymes, D9 and D10, which exhibit 25% sequence identity. Both enzymes are highly conserved with D10 homologs present in all sequenced poxviruses, and D9 homologs encoded by chordopoxviruses but not entomopoxviruses. D9 is expressed early in infection and D10 is expressed only after viral DNA replication (Baldick and Moss, 1993; Yang et al., 2010). Their temporal difference in expression and redundant roles is thought to sharpen the transition between early, intermediate and late phases of virus replication by shortening the half-life of viral mRNAs (Parrish and Moss, 2006).

D9 and D10 belong to the Nudix hydrolase superfamily of enzymes. Nudix enzymes catalyze the hydrolysis of nucleoside diphosphates linked to any moiety and contain the conserved 23-residue catalytic Nudix/MutT motif GX_5_EX_7_REUXEEXGU, where X is any amino acid and U is a bulky, hydrophobic amino acid (isoleucine, leucine, or valine) (Mildvan et al., 2005). This motif forms a loop-α helix-loop structure to function as a Mg^2+^-binding and catalytic site, which is part of a common α/β/α-sandwich or Nudix fold. Although the substrates and mechanisms of Nudix hydrolases are quite diverse, hydrolysis typically occurs by nucleophilic substitution at phosphorus and requires a varied number of divalent cations. Substrate specificity is determined by amino acid side chains outside the Nudix motif with further variation facilitated by additional insertions or domains outside the canonical Nudix fold (Mildvan et al., 2005). It has previously been demonstrated that mutation of the glutamate residues of the EUXEE sequence of the Nudix motif abolish D9 and D10 decapping activity (Parrish and Moss, 2007; Parrish et al., 2007; Soulière et al., 2010). This is similar to the eukaryotic decapping enzyme Dcp2, in which the analogous glutamate residues coordinate two Mg^2+^ ions and facilitate cap hydrolysis (She et al., 2006). Mutating these conserved glutamates in Dcp2 abolishes metal ion binding to inactivate the enzyme (Aglietti et al., 2013), which is likely the same mechanism of inactivation of D9 and D10.

In addition to the key roles poxvirus decapping enzymes play to shut off host protein synthesis and sharpen the transition between stages of the viral replication cycle, they are crucial to reduce the accumulation of viral dsRNA and therefore prevent induction of innate immune responses. Infection with vaccinia virus (VACV) containing catalytic mutations to both D9 and D10 results in a 15-fold increase in dsRNA, activation of host restriction factors protein kinase R (PKR) and 2’-5’ oligoadenylate synthetase (OAS), and inhibition of virus replication (Liu et al., 2015). Mice are able to restrict D9- or D10-deficient VACV replication in normal cells but not cancer cells, which often have impaired interferon signaling (Burgess et al., 2018; Critchley-Thorne et al., 2009). It has also recently been shown that VACV decapping enzymes are required for selective translation of viral post-replicative mRNAs (Cantu et al., 2020). The importance of D9 and D10 for viral replication suggests that these enzymes are valuable targets for antiviral therapeutics.

High resolution structural information of poxvirus decapping enzymes is critical to develop specific antiviral therapeutics to avoid off-target interactions with host decapping enzyme and cap binding proteins. Here we present the first crystal structure of VACV D9 in a product-bound inactive conformation. This structure reveals that D9 consists of two domains: a conserved Nudix domain and a three-helix bundle domain. Unlike Dcp2 in which the catalytic Nudix and regulatory domains are connected end-to-end, the three-helix bundle domain of D9 is inserted within the Nudix domain. The m^7^G cap is positioned in a composite binding site composed of residues in both the domains. Specificity for the methylated guanosine cap is achieved by stacking between conserved aromatic residues. We show that the cap is recognized during the catalytic step and not during substrate binding. We further demonstrate that D9 does not exhibit selectivity for the first transcribed nucleotide identity and is able to efficiently remove the cap from dsRNA substrates. This promiscuity with regards to substrate selection allows for the simultaneous suppression of host protein synthesis and minimization of viral dsRNA accumulation. Our structure elucidates how this decapping enzyme is able to accommodate a range of RNA substrates. The fact that cap and RNA body recognition differs from that of eukaryotic Dcp2 supports poxvirus decapping enzymes as viable antiviral targets.

## RESULTS

### Novel Domain Architecture of D9 Revealed by X-Ray Crystallography

To better understand the structural features important for D9 cap recognition, we determined the crystal structure of wild-type vaccinia virus D9 bound to its m^7^GDP product to 1.7 Å (**Table 1; Figure 1A**). This crystal structure is the first 3D structure of a poxvirus decapping enzyme and reveals a previously unpredicted 3 helix bundle domain that is inserted within the primary sequence of the Nudix domain (**Figure 1A**). This insertion domain contacts the Nudix domain through a conserved buried surface interface of 705 Å^2^. Unsurprisingly, the Nudix domain exhibits a characteristic α/β/α-sandwich fold common to all Nudix enzymes (Hua et al., 2008).

**Table 1.** Data collection and refinement statistics.

|  | D9 + m<sup>7</sup>GDP (PDB 7SEZ) | D9mu + trinucleotide (PDB 7SF0) |
| --- | --- | --- |
| **Data collection** |  |  |
| Space group | C222<sub>1</sub> | C222<sub>1</sub> |
| Cell dimensions |  |  |
| a, b, c (Å) | 52.55, 140.1, 75.98 | 52.74, 139.99, 76.02 |
| α, β, γ (°) | 90, 90, 90 | 90, 90, 90 |
| Resolution (Å) | 51.5-1.7 (1.761-1.7)<sup>a</sup> | 70-1.95 (2.02-1.95)<sup>a</sup> |
| R<sub>merge</sub> | 0.05939 (2.264) | 0.08334 (1.142) |
| I/σ(I) | 26.5 (2.81) | 20.86 (1.96) |
| Completeness (%) | 99.83 (99.58) | 99.76 (98.39) |
| Redundancy | 12.8 (12.7) | 12.6 (9.3) |
| **Refinement** |  |  |
| Resolution (Å) | 51.5-1.7 | 70-1.95 |
| No. unique reflections | 31,249 | 20,891 |
| R<sub>work</sub>/R<sub>free</sub> | 0.1935/0.2245 | 0.2010/0.2471 |
| No. atoms |  |  |
| Protein | 1,729 | 1,717 |
| Ligand/ion | 38 | 32 |
| Water | 198 | 169 |
| B-factors |  |  |
| Protein | 36.4 | 36.7 |
| Ligand/ion | 47.0 | 47.8 |
| Water | 45.1 | 42.7 |
| R.M.S. deviations |  |  |
| Bond lengths (Å) | 0.008 | 0.005 |
| Bond angles (°) | 1.22 | 0.92 |
| Ramachandran (%) |  |  |
| Favored | 96 | 97 |
| Allowed | 3.04 | 3 |
| Outliers | 0.96 | 0 |

Each dataset was collected from a single crystal.
<sup>a</sup>Values in parentheses are for the highest resolution shell.

**Figure 1.**
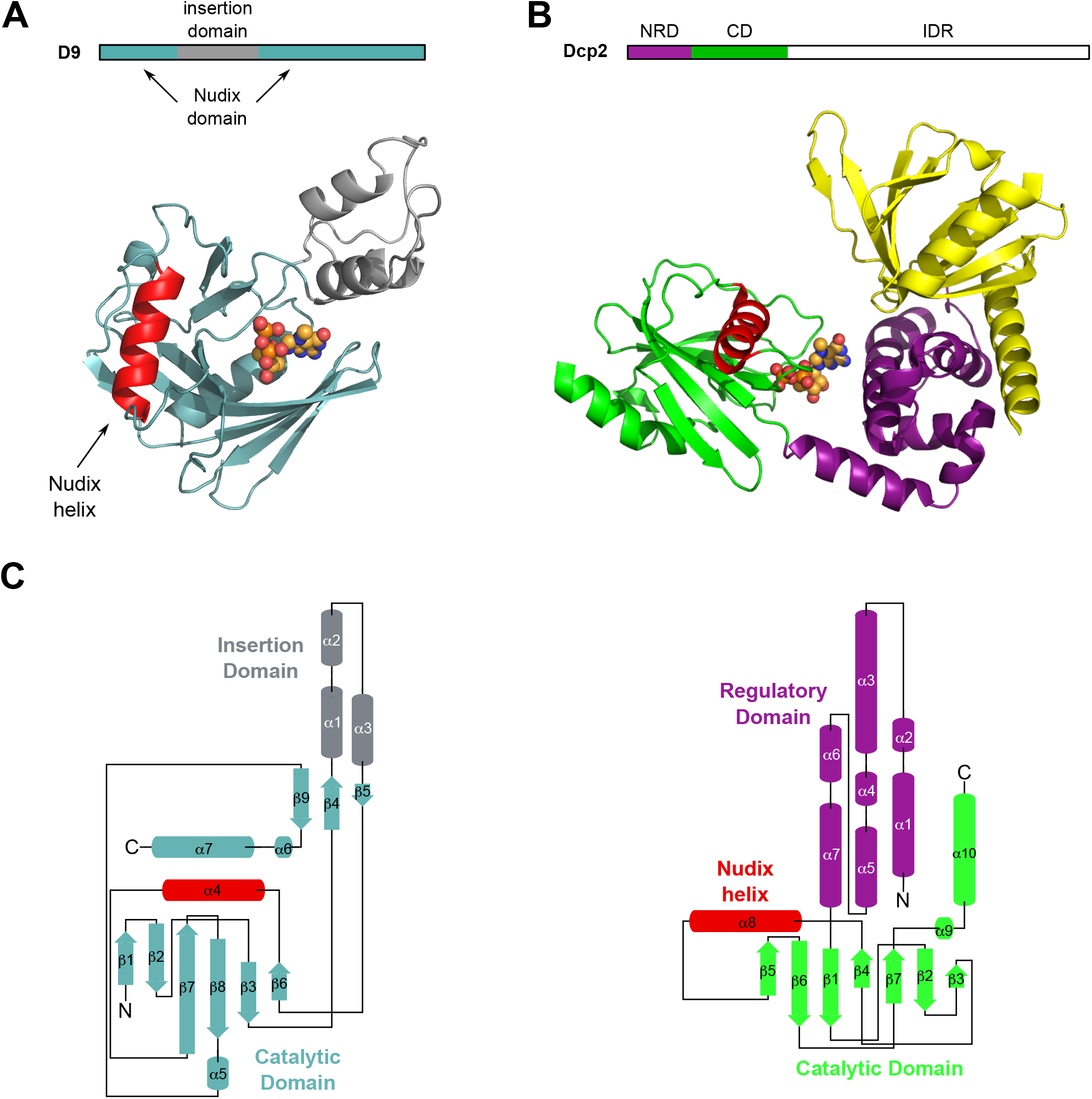
Crystal structure of poxvirus decapping enzyme D9. (A) Block diagram of VACV-WR D9 domains (above) and crystal structure of VACV-WR D9 bound to m^7^GDP product PDB 7SEV (below). D9 catalytic Nudix domain is teal, novel insertion domain is grey, Nudix helix is red, m^7^GDP is represented as spheres. (B) Block diagram of *S. pombe* Dcp2 domains (above) and crystal structure of the m^7^GDP product bound *S. pombe* Dcp1/Dcp2 decapping complex bound to m^7^GDP PDB 5N2V (below). NRD is the N-terminal regulatory domain, CD is the catalytic domain, IDR is the intrinsically disordered region. Dcp1 is yellow, Dcp2 N-terminal regulatory domain (NRD) is purple, Dcp2 catalytic Nudix domain (CD) is green, Nudix helix is red, m^7^GDP is represented as spheres. (C) Topology maps of D9 (left) and Dcp2 (right). Coloring is the same as in (A) and (B).

This is the first Nudix enzyme to our knowledge that contains a separate domain inserted within the Nudix sequence, and it is conserved among all D9 sequences (**Figure S1**). In comparison, the regulatory domain of eukaryotic decapping enzyme Dcp2 is connected to the N-terminus of its catalytic Nudix domain (**Figure 1B, 1C**). A structural homology search of the D9 insertion domain using the Dali server returned no hits. Similarly, no homologous sequences outside poxviridae were determined by the NCBI BLAST server. Therefore, the evolutionary origin of this novel 3 helix bundle domain remains unclear.

### The mRNA Cap Binds to a Composite Binding Site

The 350 Å^2^ m^7^GDP binding pocket is located at the interface of the insertion and Nudix domains where the m^7^G is positioned by stacking with conserved aromatic residues F54 and Y158 (**Figure 2A, 2B**). This mode of cap recognition is the same that is observed in canonical cap binding proteins VP39 (Hodel et al., 1997), eIF4E (Marcotrigiano et al., 1997; Matsuo et al., 1997) and CBP20 (Mazza et al., 2002) in which the m^7^G cap is sandwiched between two aromatic residues. Previous biochemical work with these proteins has demonstrated that methylated cap specificity is achieved by the favorable energetics from continuous stacking with a methylated vs. unmethylated guanine base (Hodel et al., 1997; Hsu et al., 2000; Marcotrigiano et al., 1997; Niedzwiecka et al., 2002; Quiocho et al., 2000; Stolarski et al., 1996). In our crystal structure, conserved residues D151 and E16 also form hydrogen bonds with the Watson-Crick and sugar edges of the guanine base, respectively (**Figure 2B**). Similar base pair mimicry to recognize the m^7^G cap is also employed by cap binding proteins eIF4E, CBP20, Dcp2 and DcpS (Charenton et al., 2016; Gu et al., 2004; Mazza et al., 2002; Mugridge et al., 2018a; Peter et al., 2015; Wurm et al., 2017).

**Figure 2.**
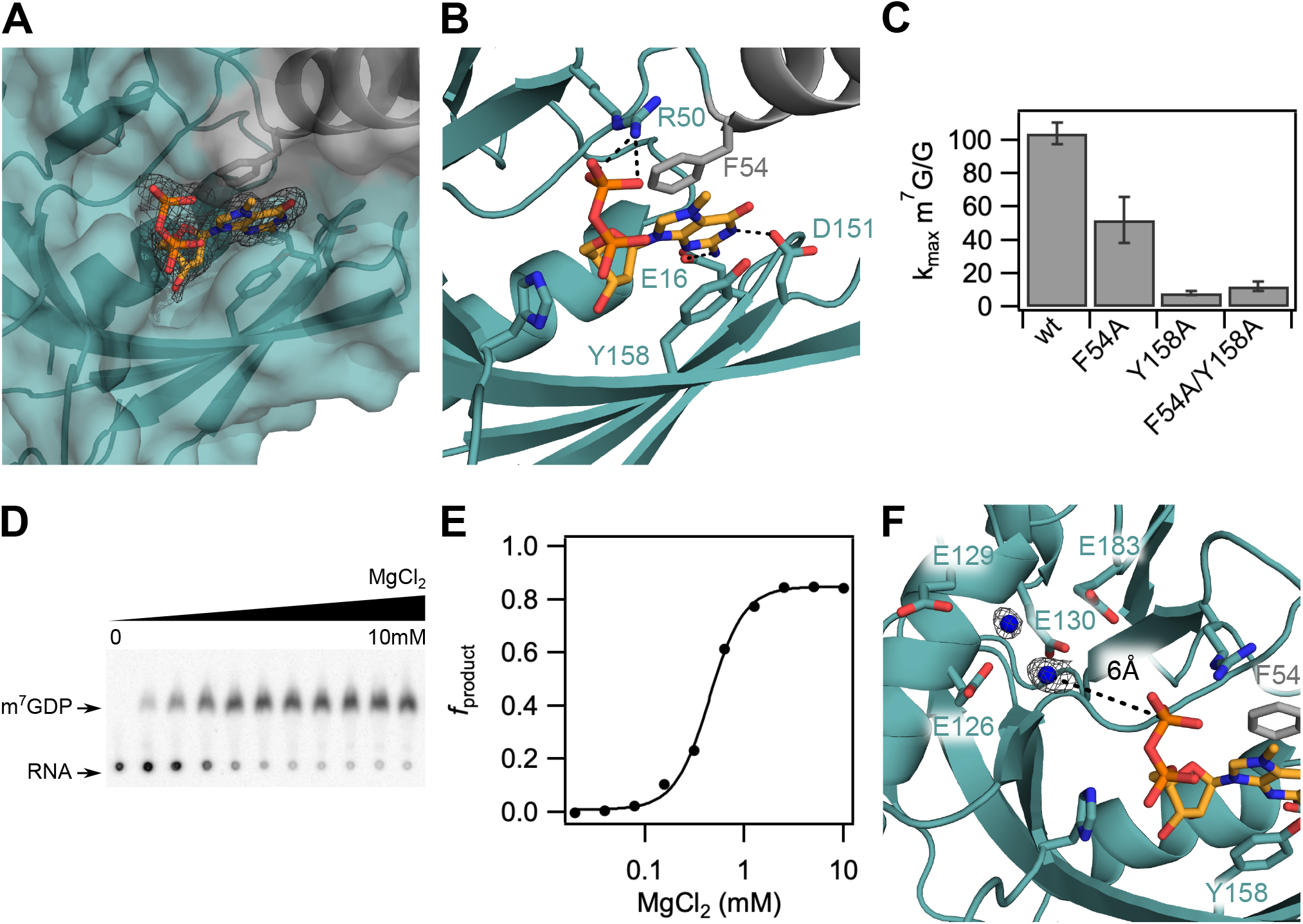
The m^7^G cap is positioned in a composite nucleotide binding site in D9. (A) Surface view of the m^7^GDP binding pocket located at the interdomain interface of the Nudix (teal) and insertion (grey) domains. F_o_-F_c_ omit map is shown for m^7^GDP substrate at 2.5σ. (B) Close up view showing interactions between the m^7^GDP substrate and both domains of D9. The methylated guanine base is sandwiched between conserved aromatic residues F54 and Y158, and hydrogen-bonds with E16 and D151 on the sugar and Watson-Crick edges, respectively. The phosphate chain is stabilized by hydrogen-bond contacts with R50. (C) Bar graph showing methylated cap specificity (ratio of decapping rates of RNA containing methylated cap to unmethylated cap) for wild-type D9 and alanine substitutions of aromatic m^7^G binding residues. Error bars are s.e.m. for the rate measured in two independent experiments. (D) TLC plate monitoring cap hydrolysis of wild-type D9 at varied [MgCl_2_] (0 to 10mM). (E) Plot of fraction m^7^GDP product of wild-type D9 vs. magnesium concentration (Hill coefficient = 2.5; [Mg^2+^]_1/2_ = 0.44mM). (F) Close up view of the conserved glutamate residues in the catalytic Nudix helix. Four conserved glutamate residues (E126, E129, E130, E183) are shown as sticks. Water molecules are shown as blue spheres. F_o_-F_c_ omit map for the waters is shown at 2.5σ in black. The distance between the β phosphate and the nearest water is 6.0 Å (black dashed line).

In principle, this binding pocket could interact with the m^7^G cap or first transcribed nucleotide. It is unclear from our crystal structure alone and comparison to available crystal structures of Dcp2 which nucleotide of the mRNA substrate occupies this position. To test the prediction that D9 uses F54 and Y158 to interact with substrate, we performed kinetic analyses of mutants under single turnover conditions where enzyme is in excess of substrate (Jones et al., 2008). F54 and Y158 are important for catalysis, as mutation of F54 or Y158 to alanine reduces the rate of the catalytic step (*k*_max_) of methylated RNA by 42- and 98-fold, respectively (**Figure S2A**). The observed decrease in activity is not due to aggregated protein because both F54A and Y158A mutant proteins produced size exclusion profiles similar to wild-type D9 (**Figure S2B**). If F54 and Y158 recognize the mRNA cap, then mutation of these residues should decrease m^7^G cap specificity. To test this hypothesis, we mutated these residues to alanine and performed decapping assays using RNAs containing a canonical m^7^G cap and noncanonical unmethylated G cap. Wild-type D9 exhibits a 100-fold reduction in *k*_max_ for unmethylated RNA compared to methylated RNA (*k*_max_ m^7^G/G = 100), indicative of specificity for methylated cap as was previously demonstrated (**Figure 2C**) (Parrish and Moss, 2007). Mutation of F54 or Y158 reduces this specificity by 2- and 13-fold, respectively, consistent with a role in cap binding. Indeed, The Y158 alanine substitution almost completely abolishes sensitivity to cap methylation (*k*_max_ m^7^G/G = 7), affirming that Y158 plays a principal role in conferring m^7^G specificity. Therefore, we conclude that the m^7^GDP substrate in our crystal structure is positioned in the cap binding pocket. Notably, the double mutant *k*_max_ m^7^G/G is equivalent to that observed for the Y158A single mutation (**Figure 2C**). This non-additive effect confirms the F54 and Y158 residues cooperatively position the methylated base through continuous π-π stacking and suggests these mutations may also change the reaction mechanism (Wells, 1990).

Previous biochemical work has shown that D10 contains two metal ion binding sites and is able to catalyze cap hydrolysis in the presence of either metal ion, with a higher activity when both are present (Soulière et al., 2009). Similarly, magnesium-dependent decapping kinetics of Dcp2 fit a Hill coefficient of 2.4, suggesting Dcp2 also binds at least two metal ions (Jones et al., 2008). As expected, analysis of D9 decapping at varied [Mg^2+^] also reveals a cooperative dependence on magnesium for cap hydrolysis with a Hill coefficient of 2.5 (**Figure 2D, 2E**). Although this Hill coefficient suggests a model in which at least two metal ions bind, we do not see any evidence for magnesium binding in our crystal structure despite the presence of magnesium in the crystallization buffer (**Figure 2F**). Though it is unknown for D9 whether cap hydrolysis proceeds through nucleophilic attack of the α or β phosphate, the closest water molecule is too far away (6 Å from the β phosphate) to proceed through either mechanism (**Figure 2F**). Therefore, we conclude that our crystal structure likely represents a post-catalytic product-bound conformation.

### Trinucleotide Substrate- and m^7^GDP Product-Bound D9 Conformations are Similar

To trap the substrate-bound, catalytically active conformation of D9 and resolve the essential metal ions, we co-crystallized D9 with a non-hydrolyzable trinucleotide substrate containing the cap and first two transcribed nucleotides. We solved this crystal structure to 1.95 Å by molecular replacement with PDB 7SEV (**Table 1; Figure 3A, 3B**). The enzyme conformation is nearly identical to our product-bound structure and both enzymes superimpose with an average all-atom 0.663 Å root-mean-square deviation (RMSD). Furthermore, density is only clearly visible for the m^7^G cap of our trinucleotide substrate (**Figure S3A**). As observed in our m^7^GDP bound structure, the m^7^G cap stacks between the conserved aromatic residues F54 and Y158 and forms hydrogen bonds with E16 and D151 on the sugar and Watson-Crick edges of the guanine base, respectively (**Figure S3A**). We can confidently place the m^7^G cap in this position rather than the first or second transcribed nucleotide due to the clear density for the N-7 methyl moiety (**Figure S3B**).

**Figure 3.**
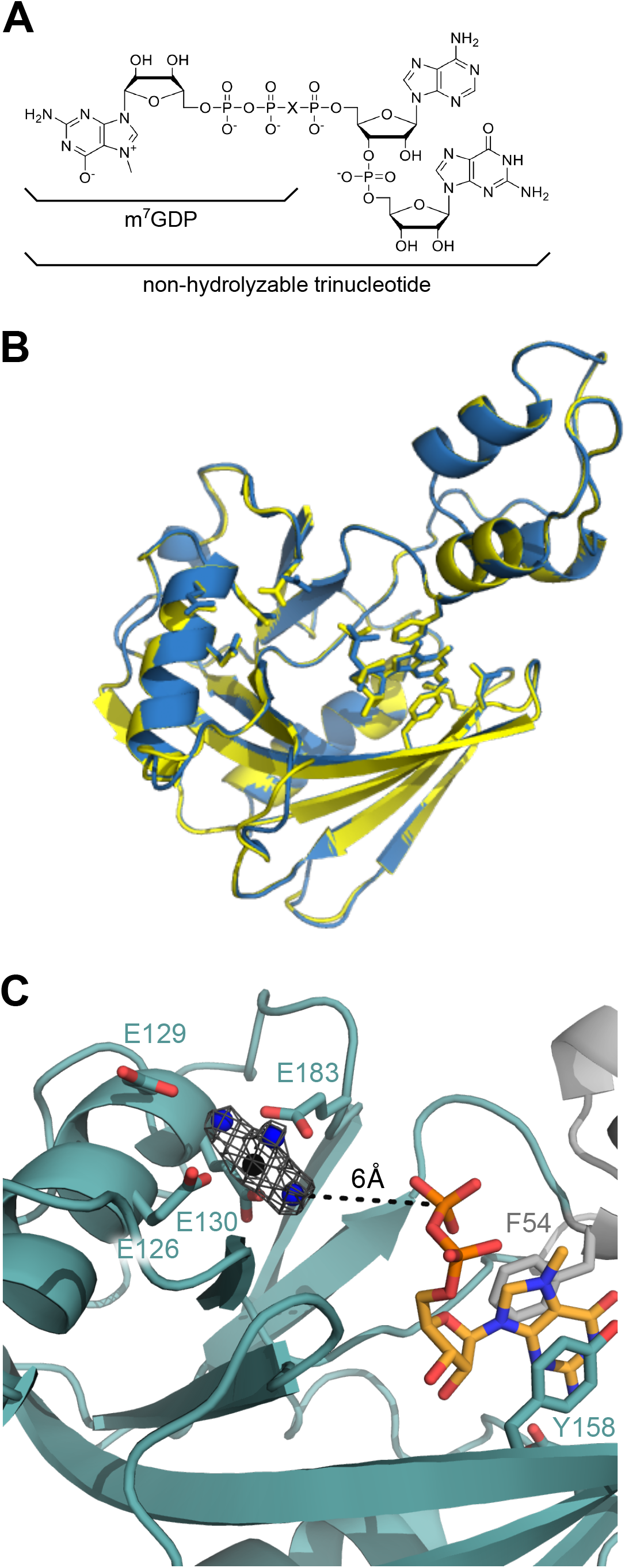
D9 likely requires RNA body to position the m^7^G for hydrolysis. (A) Molecular structures of m^7^GDP (X=O) and the trinucleotide substrate (X=CH_2_) used for co-crystallization. (B) Alignment of D9 crystal structures obtained by co-crystallizing with m^7^GDP (blue, PDB 7SEV) or trinucleotide substrate (yellow, PDB 7SF0). The backbone RMSD is 0.296 Å and all-atom RMSD is 0.663 Å. (C) Close-up of the active site and cap binding site of trinucleotide-bound D9. Four conserved glutamate residues (E126, E129, E130, E183) and residues positioning the m^7^G cap are shown as sticks. Water molecules are shown as blue spheres and the Mg^2+^ ion is shown as a black sphere. F_o_-F_c_ omit map for the waters and Mg^2+^ ion is shown at 4.0σ in gray.

In contrast to our wild-type structure, co-crystallization with the non-hydrolyzable trinucleotide substrate reveals clear density for a metal ion in the active site (**Figure 3C**). We conclude this metal ion to be magnesium because: 1) D9 enzymatic activity requires the presence of magnesium or manganese and 2) magnesium is the only metal ion included in the crystallization buffer. The three conserved glutamates in the catalytic Nudix motif coordinate the magnesium ion through both direct contact (E126, E130) and water-mediated contact (E129). An additional glutamate outside this motif also directly contacts the magnesium ion (E183). As with our product-bound structure, the nucleophilic water is again positioned too far away for catalysis to proceed rendering this conformation inactive. We conclude that a longer RNA substrate is needed to stabilize a catalytically relevant conformation of D9, which is consistent with previous endpoint assays showing that D9 requires an RNA body ≥ 12nt for efficient cap hydrolysis (Parrish and Moss, 2007).

In both D10 and Dcp2, the catalytic metal ions are coordinated by the conserved glutamate residues in the Nudix motif and mutation of these residues to glutamine abolishes metal binding and catalytic activity (Aglietti et al., 2013; Soulière et al., 2009). Mutation of the analogous glutamates to glutamine in D9 is similarly detrimental to enzyme activity (Parrish and Moss, 2007). This may be due to prevention of catalytic metal ion binding or mutation of the general base in the catalytic reaction. A model generated from biochemical characterization of VACV D10 in the presence of bivalent metals suggests that E141 could serve as the general base to initiate catalysis (Soulière et al., 2009). As D9 and D10 are closely related, we can infer the general base in D9 to be the analogous residue, E126.

### The m^7^G Cap Is Recognized During the Catalytic Step

To distinguish whether m^7^G cap specificity is conferred during substrate binding or the catalytic step, we determined the *K*_m_ and *k*_max_ for m^7^G- and G-capped RNA. As previously noted, D9 shows a 100-fold reduction in *k*_max_ for unmethylated RNA compared to methylated RNA, indicating a specificity for methylated cap (**Figure 2C**). Conversely, the *K*_m_ of methylated and unmethylated RNA were found to be equivalent (1.16 and 1.09 μM, respectively) (**Figure 4A, 4B, Table 2**). Y158 and F54 confer specificity for methylated cap and their mutation reduces D9 activity, suggesting they may be important for recognizing the cap during initial RNA binding or during the catalytic step (**Figure 2B, S2A**). We determined *k*_max_ and *K*_m_ for these mutants to disambiguate these possibilities and observe *k*_max_ is strongly decreased while *K*_m_ is relatively unchanged for either mutation (**Figure S2C**). These data suggest that cap recognition occurs during the catalytic step and not during substrate binding. In agreement with this conclusion, RNAs containing a 7-methylguanosine cap (m^7^G-RNA), guanosine cap (G-RNA), or a 5’ monophosphate (p-RNA) have equivalent binding affinities measured by a filter binding assay (**Figure 4D, Table S1**). Therefore, the presence of the 5’ cap structure, regardless of N7-methylation, does not affect substrate binding.

**Table 2.**
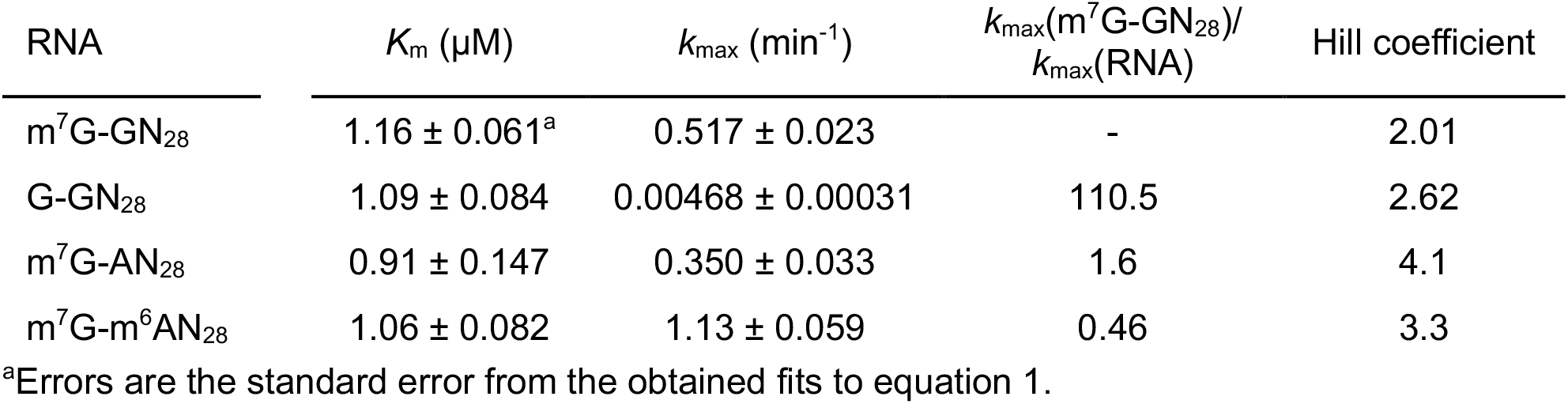
Fitted *K*_m_, *k*_max_ and Hill coefficient values for D9 decapping 29nt RNA substrates.

| RNA | $K_m$ ($\mu\text{M}$) | $k_{max}$ ($\text{min}^{-1}$) | $k_{max}(\text{m}^7\text{G-GN}_{28})/$<br>$k_{max}(\text{RNA})$ | Hill coefficient |
| --- | --- | --- | --- | --- |
| $\text{m}^7\text{G-GN}_{28}$ | $1.16 \pm 0.061^a$ | $0.517 \pm 0.023$ | - | 2.01 |
| $\text{G-GN}_{28}$ | $1.09 \pm 0.084$ | $0.00468 \pm 0.00031$ | 110.5 | 2.62 |
| $\text{m}^7\text{G-AN}_{28}$ | $0.91 \pm 0.147$ | $0.350 \pm 0.033$ | 1.6 | 4.1 |
| $\text{m}^7\text{G-m}^6\text{AN}_{28}$ | $1.06 \pm 0.082$ | $1.13 \pm 0.059$ | 0.46 | 3.3 |

<sup>a</sup>Errors are the standard error from the obtained fits to equation 1.

**Figure 4.**
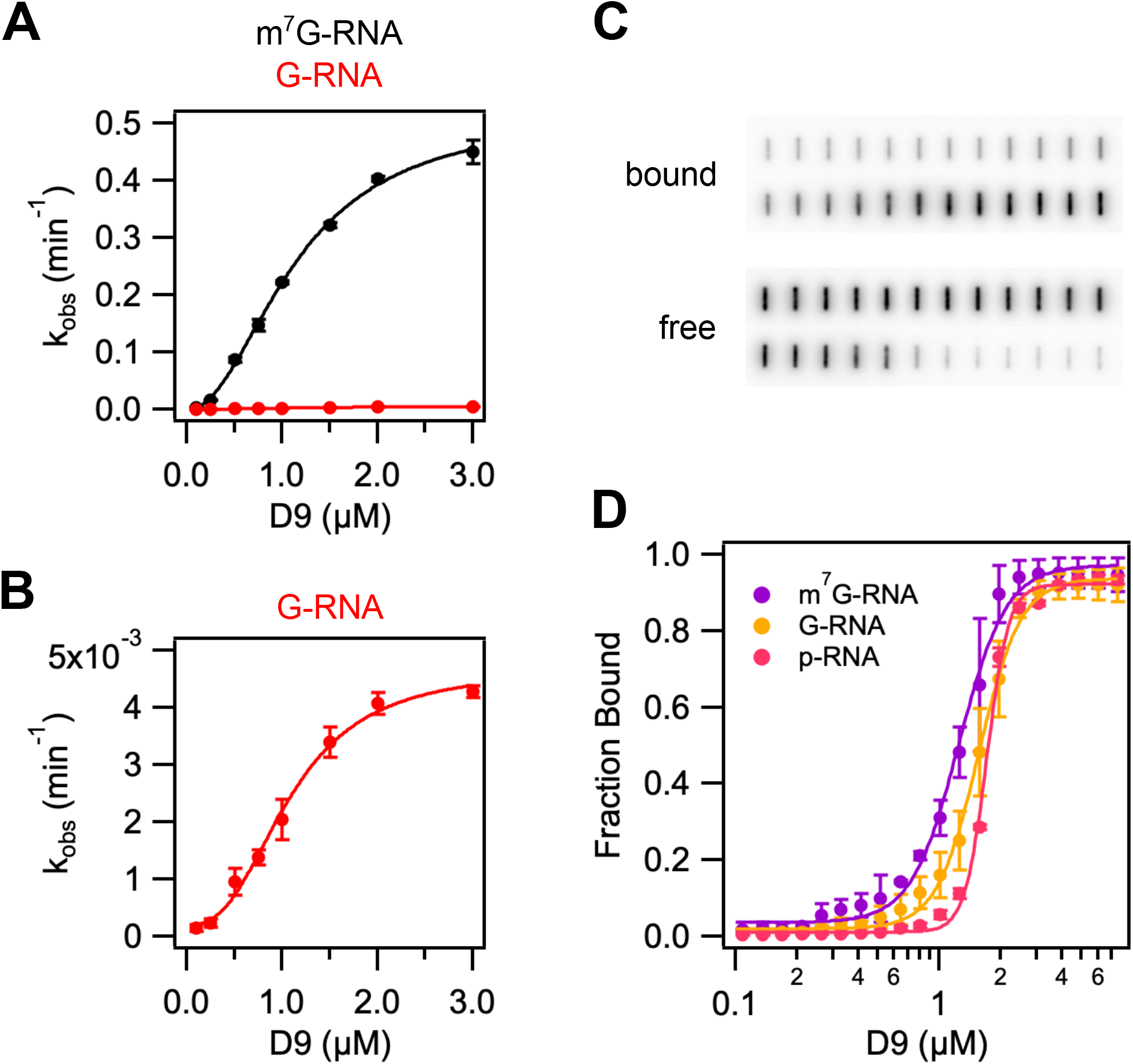
D9 recognizes methylated mRNA cap during the catalytic step by π-π stacking with conserved aromatic residues. (A) and (B) Graphs of *k*_obs_ versus wild-type D9 concentration for a 29nt RNA substrate containing a methylated (black) or unmethylated (red) guanine cap. Data were fit to Eq. 1 to determine *k*_max_, *K*_m_ and Hill coefficient (n), which are listed in Table 2. Error bars are s.e.m. for the rate measured in two independent experiments. (C) Representative filter binding assay results showing raw counts of radiolabeled RNA on nitrocellulose (bound) and Hybond N+ (free) membranes for wild-type D9 and m^7^G-capped 29nt RNA. (D) Fraction of RNA bound versus concentration of D9 for m^7^G-capped RNA (purple), G-capped RNA (yellow), and 5’ monophosphate RNA (pink). Data were fit to Eq. 3 to determine the equilibrium dissociation constants and Hill coefficients, which are shown in Table S2. Error is s.e.m. for the binding measured in two independent experiments.

The Hill coefficients for binding m^7^G-, G-, and p-RNA are 3.7, 4.2 and 8.08, respectively (**Table S1**). This is suggestive of a model in which multiple molecules of D9 assemble to bind a single RNA substrate. However, molecular weight size determination by size exclusion chromatography retention volume indicates that D9 alone is a monomeric species (**Figure S2D**). Interfaces for biological assemblies can often be observed in macromolecular crystals. A study of all protein-protein contacts in the PDB predicted 14% of these contacts are biologically relevant (Baskaran et al., 2014). Analysis of protein interfaces observed in our crystal structure using the PDBePISA and EPPIC servers did not reveal any specific interactions that could result in the formation of stable quaternary structures (Bliven et al., 2018; Krissinel and Henrick, 2007). We therefore conclude that D9 may require the RNA body or an alternative conformation to form stable assemblies.

The fact that the substrate *K*_m_ is similar to the *K*_d_ for m^7^G capped and 5’ monophosphate RNA implies the enzyme substrate falls apart rapidly compared to the rate of the catalytic step (*k*_off_ >> *k*_max_). These findings suggest that the catalytic step is the rate limiting step for decapping. Interestingly, when Y158 is mutated to alanine, D9 cannot interact with 5’-monophosphate RNA (**Figure S2E**). This effect is specific to Y158 as the equivalent mutation at F54 causes a modest defect in binding 5’ monophosphate RNA. This result indicates Y158 has a role in both RNA binding and catalysis. These enzymatic properties are similar to Dcp2, which also recognizes the m^7^G cap during the rate limiting catalytic step and contains multifunctional residues (Deshmukh et al., 2008).

### D9 does not exhibit selectivity for the first nucleotide identity

Both mammalian and poxvirus transcripts preferentially contain a purine at the first transcribed nucleotide position (Boone and Moss, 1977; Carninci et al., 2006; Tamarkin-Ben-Harush et al., 2017). However, early and late poxvirus transcripts differ in the first transcribed nucleotide identity. Unlike early transcripts which can contain either guanine or adenosine at the 5’ terminal position, the conserved TAAAT promoter sequence of VACV intermediate and late transcripts results in a 5’ poly(A) leader of variable length at the very 5’ end of the mRNA (Baldick and Moss, 1993; Davison and Moss, 1989; de Magistris and Stunnenberg, 1988; Schwer et al., 1987). VACV mRNAs beginning with adenosine increase from 37% present early in infection to 90% late in infection (Boone and Moss, 1977). These adenosine residues can also be modified by N6-methylation (m^6^A), though the ratio of A to m^6^A at the 5’ terminal position does not significantly change from early to late in infection (Boone and Moss, 1977).

As the 5’ terminal sequences of VACV mRNAs vary throughout infection, we asked whether D9 shows selectivity for the first transcribed nucleotide identity. We compared binding affinities and decapping kinetics for RNA substrates where the first transcribed nucleotide was either guanosine (G-RNA), adenosine (A-RNA), or N^6^-methyladenosine (m^6^A-RNA) (**Table 2, Figure 5A**). Binding affinities and *K*_m_ were found to be comparable for all substrates. Although we observe a two-fold increase in *k*_max_ for m^6^A-RNA compared to G-RNA, this difference is within the variability in *k*_max_ observed from different preparations of radiolabeled RNA (**Figure S4**). Therefore, we conclude that D9 shows no selectivity for G versus A or m^6^A at the first transcribed nucleotide position and consequently is capable of removing the 5’ cap from host and viral transcripts present throughout infection.

**Figure 5.**
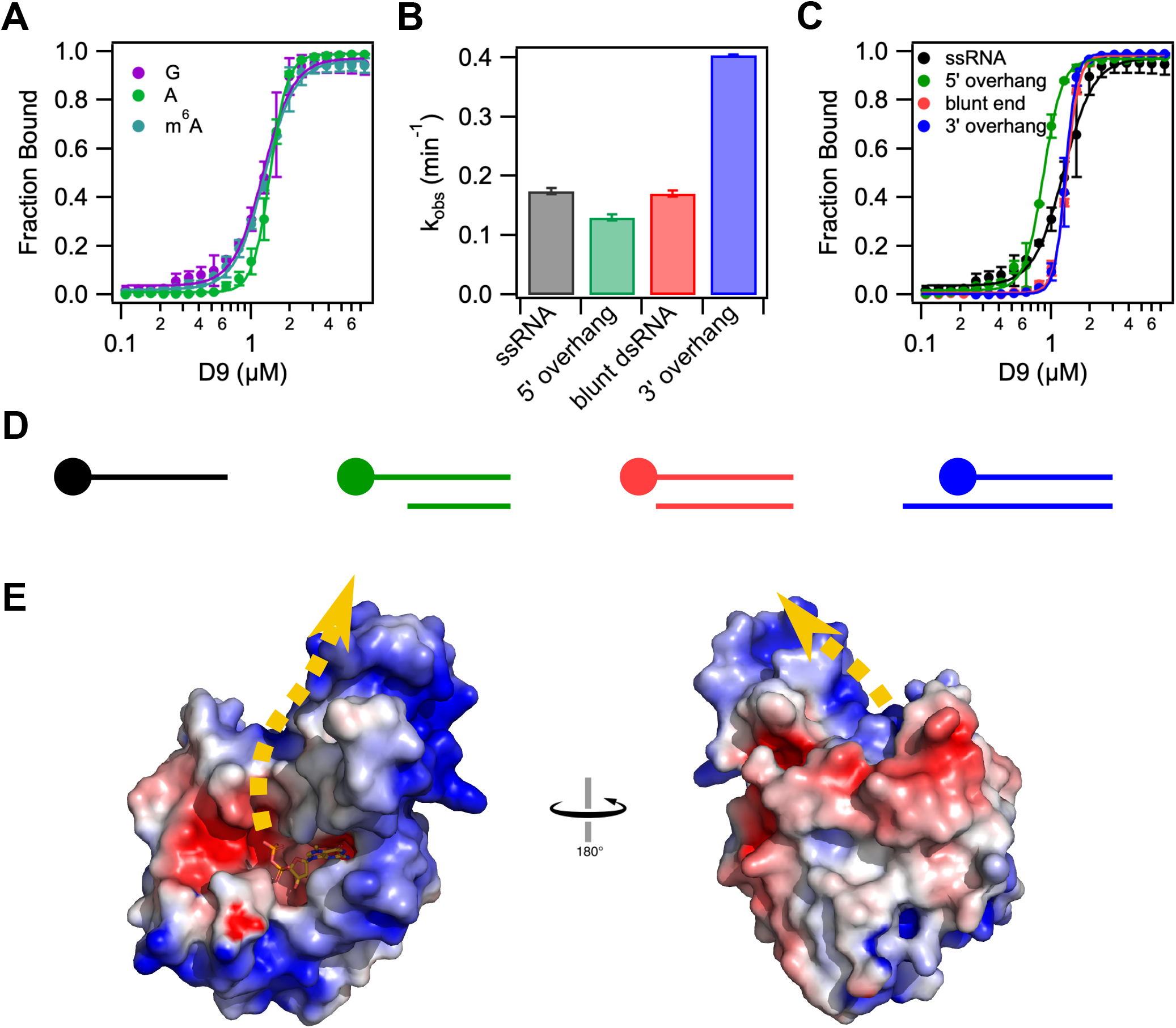
D9 does not show selectivity for first transcribed nucleotide identity or cap accessibility. (A) Fraction of RNA bound versus concentration of D9 for RNAs containing a guanosine (purple), adenosine (green) or N6-methyl adenosine (teal) at the first transcribed nucleotide position. (B) Bar graph showing the decapping rates of single stranded and double-stranded RNA constructs. All constructs contain the same m^7^G-capped 29nt RNA. Double stranded RNA constructs are annealed to complementary RNA of variable length to engineer a range of cap accessibility from fully accessible (10nt 5’ overhang) to “inaccessible” (10nt 3’ overhang). (C) Fraction of RNA bound versus concentration of D9 for the same RNA constructs used in (B). The equilibrium dissociation constants and Hill coefficients for all equilibrium binding assays are shown in Table S2. (D) Schematic representation of dsRNA constructs used in (B) and (C) from most accessible (left) to least accessible m^7^G cap (right). The m^7^G cap is shown as a circle on the top strand. Anti-sense RNA sequences are listed in Table S2. (E) Electrostatic surface potential of product-bound D9 (- 5 to +5 kT/e). Error bars show the s.e.m for the binding or rate measured in two independent experiments. Dashed yellow line represents the predicted RNA path based on the orientation of the cap and location of positively charged patches on the protein surface.

It has been shown that D9 and D10 are important to prevent accumulation of dsRNA during VACV infection. Catalytic mutations to both enzymes severely attenuates the virus, resulting in a 15-fold increase in dsRNA and induction of interferon alpha and beta (Liu et al., 2015). These enzymes could function by degrading RNAs prior to annealing or by degrading dsRNA. To determine if D9 is capable of utilizing dsRNA as a substrate, we performed decapping assays on dsRNA constructs with varied cap accessibility. Complementary RNAs were annealed to radiolabeled m^7^G-RNA to produce constructs in which the m^7^G cap ranges from fully accessible (5’ overhang) to less accessible (blunt end, 3’ overhang) (**Figure 5D**). Interestingly, the 5’ overhang and blunt end dsRNA constructs are decapped at the same rate as ssRNA, and the dsRNA construct with the least accessible cap (3’ overhang), shows a two-fold increase in *k*_max_ (**Figure 5B**). Binding affinities for all dsRNA constructs are nearly identical to the binding affinity of ssRNA, though cooperativity increases with decreasing cap accessibility (**Figure 5C, Table S1**). Therefore, D9 exhibits no preference for ssRNA versus dsRNA.

Nucleic acid binding proteins often have large positive patches on their surfaces to bind their electrostatically negative substrates in a manner independent of nucleic acid sequence. Such is the case in Dcp2 which contains a conserved positive patch on the surface of the Nudix domain (Box B) known to be critical for RNA binding (Deshmukh et al., 2008; Piccirillo et al., 2003; She et al., 2008). Though sequence alignment with Dcp2 does not reveal a homologous Box B motif in D9, we hypothesize that the RNA body likely binds in a similar way to an accessible positive patch on the surface of D9 in order to accommodate both ss- and dsRNA substrates in a non-sequence specific manner. The three-helix bundle insertion domain contains a positive patch on the posterior face of D9 from the cap binding site (**Figure 5E**). Although the identity of the residues in this region are not conserved (**Figure S1**), the positive electrostatic surface potential is conserved among nearly all chordopoxviruses (**Figure S5**). If we assume that the path of the electrostatically negative RNA bound to D9 will follow the track of the positive potential, the most likely path extends up from the cap binding pocket along the interface of the two domains and onto the three-helix bundle domain (**Figure 5E**). While the trinucleotide substrate used for co-crystallization is too short to contact the positive patch on the surface of the three-helix domain, our model of substrate binding is in agreement with the requirement for longer RNA substrate for efficient decapping (**Figure 3A**) (Parrish and Moss, 2007).

## DISCUSSION

We solved the first crystal structure of the D9 poxvirus decapping enzyme in a post-catalytic state and uncovered its catalytic mechanism. D9 contains a previously unannotated three helix bundle inserted into a highly conserved Nudix domain. Together, these domains form a bipartite active site that sandwiches the m^7^G cap between two aromatic residues (F54 and Y158). Mutating either of these residues abrogates catalysis and results in a loss of D9 specificity for N7-methylation, consistent with these interactions being important for recognizing m^7^G-capped RNA. D9 does not have specificity for the first-transcribed nucleotide and can decap ssRNA and dsRNA equally. These results indicate D9 can engage with a wide array of capped host and viral RNA to promote their degradation and ensure viral pathogenesis.

Our crystal structure confirms D9 is similar to other Nudix hydrolases, which contain a conserved catalytic helix and additional domains that confer specific substrate recognition (Mildvan et al., 2005). D9 has a novel domain inserted within the Nudix fold that has residues important for stabilizing m^7^GDP in a composite active site (**Figure 1A**). This domain architecture is most similar to the eukaryotic mRNA decapping enzyme Dcp2, which recognizes m^7^G at the interface between regulatory and catalytic domains (**Figure 1B**). A flexible linker connects the regulatory and catalytic domains of Dcp2, affording much conformational plasticity important for its catalytic cycle (Wurm and Sprangers, 2019). Similarly, two loops connect the insertion domain to the Nudix domain of D9 and we predict the insertion domain can also undergo conformational changes to position substrate in the active site because the β-phosphate is positioned too far from the catalytic helix in the current structures (**Figures 2F, 3C**). High-resolution studies of the active state of D9 will be required to elucidate the extent of these conformational changes.

D9 exhibits 100-fold specificity for the N7-methylation of capped RNA substrate. The methylated guanosine base is stacked between conserved aromatic residues F54 of the insertion domain and Y158 of the Nudix domain. Additional hydrogen-bonding interactions between D9 and the guanosine Watson-Crick face reinforce this orientation. The continuous π-cation-π stacking and Watson-Crick mimicry observed are conserved mechanisms for conferring specific m^7^G recognition and is employed by several cap binding proteins including vaccinia virus RNA 2’-O-methyltransferase VP39 (Hodel et al., 1997), cytoplasmic cap-binding protein eIF4E (Marcotrigiano et al., 1997; Matsuo et al., 1997) and nuclear cap-binding complex subunit CBP20 (Mazza et al., 2002). Mutation of Y158 in the Nudix domain abolishes specificity for methylated, capped RNA. This mode of recognition is in contrast to Dcp2, wherein specificity is encoded by π-stacking with a single aromatic residue (W43) and Watson-Crick mimicry (D47) in the regulatory domain (Floor et al., 2010). The additional cation-π stacking likely stabilizes D9 interaction with m^7^G and may serve to enhance the basal decapping rate since the cap is recognized during the catalytic step and is rate-limiting.

We show D9 does not exhibit a significant preference for the first transcribed nucleotide identity and can decap ssRNA and dsRNA with similar activity (**Figure 5, Table S1**). This is in contrast to Dcp2, which prefers a purine base as the first-transcribed nucleotide and has been shown to be less active on transcripts containing m^6^A at this position (Mauer et al., 2017; Mugridge et al., 2018a). While our structures do not contain additional base pairs 3’ to the cap, we observe a large basic surface on D9 that likely acts as an RNA binding interface. This extensive positively-charged region would engage with multiple nucleotides in the mRNA body and accommodate different binding poses, explaining why D9 requires RNA substrates ≥12nt for efficient catalysis and can decap an expanded substrate profile. Altogether, these results support the role of D9 in non-discriminately decapping cellular and viral mRNAs to simultaneously: 1) shut off host protein synthesis, 2) sharpen the transition between stages of the viral replication cycle, and 3) reduce the accumulation of viral dsRNA both before and after their formation.

Efficient mRNA decapping in eukaryotes involves the assembly of large decapping complexes that enhance Dcp2 activity and deliver substrates to Dcp2 (Mugridge et al., 2018b). This coordinated regulation ensures specific transcripts are targeted for degradation in regulated mRNA decay and quality control pathways. The interactions regulating Dcp2 decapping are mainly mediated through short linear interaction motifs (SLiMs) found in a C-terminal disordered region and removal of this region leads to non-specific mRNA degradation. Poxvirus D9 lacks a disordered region and our structural data suggests it most closely resembles core subunits of Dcp2. As a result, we predict D9 would not co-opt regulators of Dcp2 but instead target host or viral transcripts through different mechanisms. Our biochemical data suggests D9 can cooperatively assemble into oligomeric complexes on RNA substrate and such structures may represent degradation centers during infection. Additionally, the Dcp2 regulatory domain forms a complex with Dcp1 to collectively activate decapping 1000-fold and it is possible the insertion domain in D9 can also complex with host or viral factors to enhance activity. Assessing D9 cellular localization and interacting partners during poxvirus infection is an exciting area of future research.

The fact that D9 and D10 share 25% sequence identity and have similar secondary structure predictions based on primary sequence suggests that D10 similarly contains an insertion domain, strengthening support that D9 and D10 have a common origin. Perhaps D9 resulted from a gene duplication after the divergence of chordo- and entomopoxviruses. Although D9 and D10 catalyze the same cap hydrolysis reaction to release m^7^GDP and a 5’ monophosphate RNA, D9 is more sensitive to inhibition by uncapped RNA and D10 is more sensitive to inhibition by methylated nucleotides. These subtle differences may lead to more significant variation in substrate selectivity and roles in poxvirus infection. It was recently shown that D10 but not D9 enhances translation in the absence of VACV infection with a preference for mRNAs containing a 5’-poly(A) leader (Cantu et al., 2020). Future biochemical and biophysical studies of the differences between D9 and D10 are needed to explain their divergent roles in poxvirus infection.

Disruption of host protein synthesis and the clearance of viral dsRNA are two important mechanisms for viral pathogenesis. We demonstrate how poxvirus can utilize a broad-spectrum decapping enzyme to carry out these functions. Our structural and biochemical studies can serve as a framework for exploiting differences in catalysis and substrate recognition to selectively perturb poxvirus decapping. These mechanistic insights will be important for understanding how poxvirus can effectively evade host immune responses and aid in the development of new antivirals.

## ACKNOWLEDGEMENTS

We thank J. Holton and G. Meigs at Lawrence Berkeley National Lab, Advanced Light Source beamline 8.3.1 for help with X-ray data collection and processing. This work was supported by the US National Institutes of Health (R01 2R01GM078360 to J.D.G, NRSA fellowship F32 Grant 5F32GM133084 to J.K.P.). Beamline 8.3.1 at the Advanced Light Source is operated by the University of California Office of the President, Multicampus Research Programs and Initiatives grant MR-15-328599, the National Institutes of Health (R01 GM124149 and P30 GM124169), Plexxikon Inc. and the Integrated Diffraction Analysis Technologies program of the US Department of Energy Office of Biological and Environmental Research. The Advanced Light Source is supported by the US Department of Energy under contract no. DE-AC02-05CH11231.

## AUTHOR CONTRIBUTIONS

Conceptualization, J.K.P. and J.D.G.; Methodology, J.K.P., R.W.T., and J.D.G.; Formal Analysis, J.K.P.; Investigation, J.K.P and R.W.T.; Writing – Original Draft, J.K.P. and J.D.G.; Writing – Review & Editing, J.K.P, R.W.T. and J.D.G.; Funding Acquisition, J.K.P. and J.D.G.; Resources, M.W. and J.J.; Visualization, J.K.P.; Supervision, J.J., J.D.G

## EXPERIMENTAL PROCEDURES

### Protein Expression and Purification

*E. coli* optimized VACV strain Western Reserve D9 DNA sequence was obtained as a gblock from Integrated DNA Technologies and cloned into the pET28b vector with a C-terminal hexahistidine affinity tag. This was used to transform *E. coli* BL21 DE3 (New England Biolabs), and cells were grown in LB media to OD ∼0.6. Recombinant protein expression was induced with 0.3 mM IPTG for 16h at 18C. Cells were harvested at 5,000 x g, lysed by sonication, and clarified at 20,000 x g in lysis buffer (50 mM HEPES pH 7, 500 mM NaCl, 20 mM imidazole, 10% glycerol, 10 mM 2-mercaptoethanol, 1 mM PMSF, 1 μg/ml leupeptin, 1 μg/ml pepstatin A). Recombinant D9 was purified sequentially by a Ni-NTA column followed by a heparin column (GE Healthcare) and a GE Superdex 75 16/60 gel filtration column to separate aggregates and exchange into crystallization buffer (10 mM MES pH 5.6, 300 mM NaCl, 1 mM DTT). The purified protein was concentrated to 10 mg/ml and flash frozen in liquid nitrogen. Mutants were generated by site-directed mutagenesis and purified following the same protocol. Wild-type D9 was also expressed in its selenomethionine (Se-Met) derivatized form as previously described and purified as outlined above following induction of protein expression (Doublié, 2007).

### X-Ray Crystallography

Cap analog substrate, m^7^GDP, or trinucleotide substrate was dissolved in water at a concentration of 50mM and the pH was adjusted to ∼7. Protein-substrate complex was prepared by mixing D9 and cap analog substrate in crystallization buffer containing 20 mM MgCl_2_ to a final concentration of 10mg/ml D9 and 3 mM cap analog substrate and incubated at room temperature for 30 min. Protein-substrate complex was mixed with well solution at a 1:1 ratio and crystals were grown by hanging drop vapor diffusion at room temperature. D9-m^7^GDP well solution contained 300 mM sodium citrate, 16% PEG 3,350. D9-trinucleotide well solution contained 284 mM sodium sulfate, 17% PEG 3,350, 100 mM Bis Tris propane pH 7.5. Crystals were flash frozen in liquid nitrogen using a cryoprotectant consisting of well solution with 25% glycerol. All data sets were collected on beamline 8.3.1 at the Advanced Light Source at 100K and either 0.9795 Å (Se SAD phasing) or 1.115869 Å (native datasets) using the Pilatus3 S 6M detector and indexed, integrated and scaled using XDS (Kabsch and IUCr, 2010), Pointless and Aimless (Winn et al., 2011) via automated beamline software ELVES (Holton and Alber, 2004). Phases were determined by Se SAD phasing of data sets collected at 0.9795 Å from a single SeMet-D9-m^7^GDP crystal using Phenix AutoSol (Terwilliger et al., 2009). The structure was then iteratively refined in PHENIX (Adams et al., 2010) and manually adjusted in COOT (Emsley et al., 2010). The structure of D9 co-crystallized with trinucleotide substrate (PDB 7SF0) was solved by molecular replacement using PDB 7SEV.

### Accession Codes

The coordinates and structure-factor amplitudes have been deposited in the Protein Data Bank under accession codes 7SEV (m^7^GDP-bound) and 7SF0 (trinucleotide-bound).

### Synthesis of RNA Substrates

The 5’-triphosphorylated 29-nt RNAs used for capping reactions were synthesized by Trilink. The 22-, 29-, 32- and 42-nt transcripts used for preparation of double stranded RNA were synthesized by *in vitro* transcription using annealed oligonucleotide primers containing the T7 promoter sequence and the corresponding AS sequence of the 29-nt RNA with C2’-methoxyls at the last two nt of the template DNA 5’ termini to reduce nontemplated nucleotide addition (Kao et al.). RNA sequences are shown in Table S2. RNA was 5’ cap radiolabelled as previously described to generate cap 0 structures (Deshmukh et al., 2008). To prepare radiolabeled 5’ monophosphate RNA, 100pmol triphosphorylated RNA was first dephosphorylated using shrimp alkaline phosphatase (New England Biolabs) following the manufacturer’s instructions. The RNA was then phosphorylated using T4 PNK (New England Biolabs) and γ^32^P-ATP following the manufacturer’s instructions. The radiolabeled RNA was purified by gel filtration.

### Kinetic Decapping Assays

Single-turnover *in vitro* decapping assays were carried out as previously described (Jones et al., 2008) at room temperature in 10mM HEPES pH 7.5, 100mM KOAc, 1mM MgCl_2_ and 2mM DTT. Three different preparations of 29-nt RNA (denoted a, b and c) were used during these experiments. Comparison of *K*_m_ and *k*_max_ from D9wt and m^7^G-GRNA shows no significant difference in *K*_m_ and up to a 3.2-fold difference in *k*_max_ (Figure S4). Importantly, comparisons between kinetic constants are only drawn from RNAs prepared on the same day. To obtain *k*_max_, *K*_m_ and Hill coefficient (n), *k*_obs_ was plotted versus protein concentration and fitted to the model:

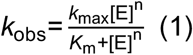

### Magnesium dependent activity assay

Single-turnover *in vitro* decapping assays were carried out as previously described (Jones et al., 2008) with 2μM D9wt in 10mM HEPES pH 7.5, 100mM KOAc, 2mM DTT, and MgCl_2_ ranging from 0 to 10mM in a 10μl reaction volume. Reactions were incubated at room temperature for 30min and quenched with 2μl 0.5M EDTA. The fraction of m^7^GDP product was measured for each [MgCl_2_] and the resulting data was fit to a standard Hill equation:

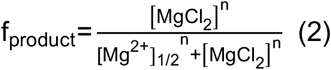

where n is the apparent Hill coefficient and [Mg^2+^]_1/2_ is the apparent magnesium concentration required for decapping one-half of the RNA substrate.

### Filter Binding Assays

Filter binding assays were carried out as previously described (Rio, 2012). Briefly, radiolabeled RNAs were incubated at room temperature with serial dilutions of D9mu in binding buffer containing 10mM HEPES pH 7.5, 100mM KOAc, 1mM CaCl_2_ and 2mM DTT for 30min. D9mu contains two point mutations (E129Q/E130Q) which have been shown to render the enzyme catalytically inactive (Parrish and Moss, 2007). The final concentration of radiolabeled RNA in each sample was <10nM. Binding reactions were loaded onto a slot blot manifold pre-washed with binding buffer and applied sequentially to nitrocellulose and Hybond N+ membranes using a vacuum to capture bound and free RNA, respectively. Dry membranes were exposed to phosphorimaging screens (Amersham) overnight and scanned with a PhosphorImager (Typhoon, GE Amersham). The fraction of RNA bound was measured for each concentration of D9 and the resulting data was fit to a standard Hill equation:

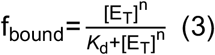

where n is the apparent Hill coefficient, *K*_d_ is the equilibrium dissociation constant and [E_T_] is the total concentration of D9.

### Structural Analysis and Visualization

Buried surface area in protein interfaces was calculated using the PDBePISA webserver (http://www.ebi.ac.uk/pdbe/prot_int/pistart.html). Electrostatic surfaces were calculated using PDB2PQR & APBS webservers (https://server.poissonboltzmann.org/) (http://nbcr-222.ucsd.edu/pdb2pqr_2.0.0/) using the AMBER forcefield to calculate charges (Baker et al., 2001; Dolinsky et al., 2004). Electrostatic surfaces were visualized in PyMol using APBS tools 2.1. Topology maps were generated using PDBsum (Laskowski et al., 2018).

### Sequence Alignments

Sequences were obtained from UniProt (Consortium et al., 2021), aligned with MUSCLE (Madeira et al., 2019) and visualized using Jalview (Waterhouse et al., 2009). Consensus sequence images were generated using Weblogo (Crooks et al., 2004).

