## Supplemental Materials for "Structure of the poxvirus decapping enzyme D9 reveals its mechanism of cap recognition and catalysis"

**Supplementary Materials**

| **Supplemental Figures** | **Page** |
| --- | --- |
| **Figure S1**: Sequence conservation of D9 among chordopoxvirinae | **3** |
| **Figure S2**: Residue Y158 plays multiple roles in substrate binding and catalysis | **4** |
| **Figure S3**: Trinucleotide substrate positioning in cap binding pocket | **5** |
| **Figure S4**: Measured decapping rate is variable between different RNA preparations | **6** |
| **Figure S5**: D9 electrostatic surface comparison of representative poxviruses | **7** |
| **Supplementary Data Tables** | |
| **Table S1**: Substrate binding affinities measured by filter binding assay | **8** |
| **Table S2**: RNA sequences used for activity and binding assays | **8** |

**Figure S1**. Sequence conservation of D9 among chordopoxvirinae. (A) Sequence alignment of representative D9 sequences. VACV-WR, vaccinia virus western reserve strain; YMTV, yaba monkey tumor virus; MYXV, myxoma virus; DPV, deerpox virus; SWPV, swinepox virus; SHPV, sheeppox virus; MCV, molluscum contagiosum virus; FWPV, fowlpox virus; CDPV, crocodylidpox virus. Background color is coded by ConSurf conservation score from variable (teal) to conserved (magenta) from multiple sequence alignment of all available D9 sequences. Numbering above the sequences follows VACV-WR residue number. Secondary structure for VACV is shown above the sequences where arrows are beta sheets and cylinders are alpha helices. Coloring is the same as Figure 1. Triangles below indicate conserved residues in the Nudix motif, consisting of the sequence $GX_5EX_7REUXEEXGU$, where X represents any residue and U represents an isoleucine, leucine or valine. (B) 3D structure of product-bound D9 with D9 shown as space-filled model and $m^7$GDP shown as sticks. Residues are colored by conservation as in (A).

**Figure S2**. Residue Y158 plays multiple roles in substrate binding and catalysis. (A) Bar graph showing the rate of the catalytic step ($k_{max}$) plotted on a log scale for cap binding mutants using methylated (blue) and unmethylated (purple) RNA substrate. Error is s.e.m. for the rate measured in two independent experiments. (B) Preparative size exclusion chromatography (SEC) of wild-type D9 (blue) and cap binding mutants F54A (yellow) and Y158A (purple) from the final step in their individual purifications (superdex 75 16/600 gel filtration column). Elution volumes are 66.2, 70.4, and 68.9ml, respectively. (C) Graphs of $k_{obs}$ versus mutant D9 concentration for decapping of $m^7$G-RNA. Each data point is from a single replicate. F54A: $K_m$=0.5μM, $k_{max}$=0.015min-1, n=2.7; Y158A: $K_m$=1.82μM, $k_{max}$=0.0048min-1, n=4.6. (D) Analytical SEC of wild-type D9 and molecular weight standards aprotinin (6.5 kDa), ribonuclease A (13.7 kDa), carbonic anhydrase (29 kDa), ovalbumin (44 kDa), and conalbumin (75 kDa). The apparent molecular weight of wild-type D9 determined from elution volume comparison to standards is 21.1 kDa. (E) Fraction of RNA bound versus concentration of D9 for $m^7$G-capped RNA (purple) and 5' monophosphate RNA (pink) for cap binding mutants F54A, Y158A and F54A/Y158A. The equilibrium dissociation constants and Hill coefficients are shown in Table S2. Error is s.e.m. for the binding measured in two independent experiments.

**Figure S3**. Trinucleotide substrate positioning in cap binding pocket. (A) Crystal structure of VACV-WR D9 bound to non-hydrolyzable trinucleotide substrate (PDB 7SF0). $F_o$-$F_c$ omit map is shown for trinucleotide substrate at 2.0σ. Only clear density is observed for the $m^7$G cap and two phosphate groups. (B) Close up view showing positioning of the $m^7$G cap between aromatic residues F54 and Y158. $F_o$-$F_c$ omit map is shown for the N7-methyl moiety at 3.5σ.

**Figure S4**. Measured decapping rate is variable between different RNA preparations. (A) Graph of $k_{obs}$ versus wild-type D9 concentration for three different preparations of $m^7$G capped 29nt RNA substrate. (B) Fitted $K_M$, $k_{max}$ and Hill coefficient values for data in (A). Errors are the standard error from the obtained fits to equation 1.

**Figure S5**. D9 electrostatic surface comparison of representative poxviruses. Poxvirus sequences from eight different genera were modelled onto the D9 crystal structure (PDB 7SEV) using MODELLER (Šali and Blundell, 1993) using sequence alignments generated using MUSCLE (Madeira et al., 2019). Electrostatic surfaces shown are -5 to +5 kT/e. Top row is sequence conservation as determined by Consurf server using multiple sequence alignment of all known D9 sequences (Figure S1).

#### Figure S1

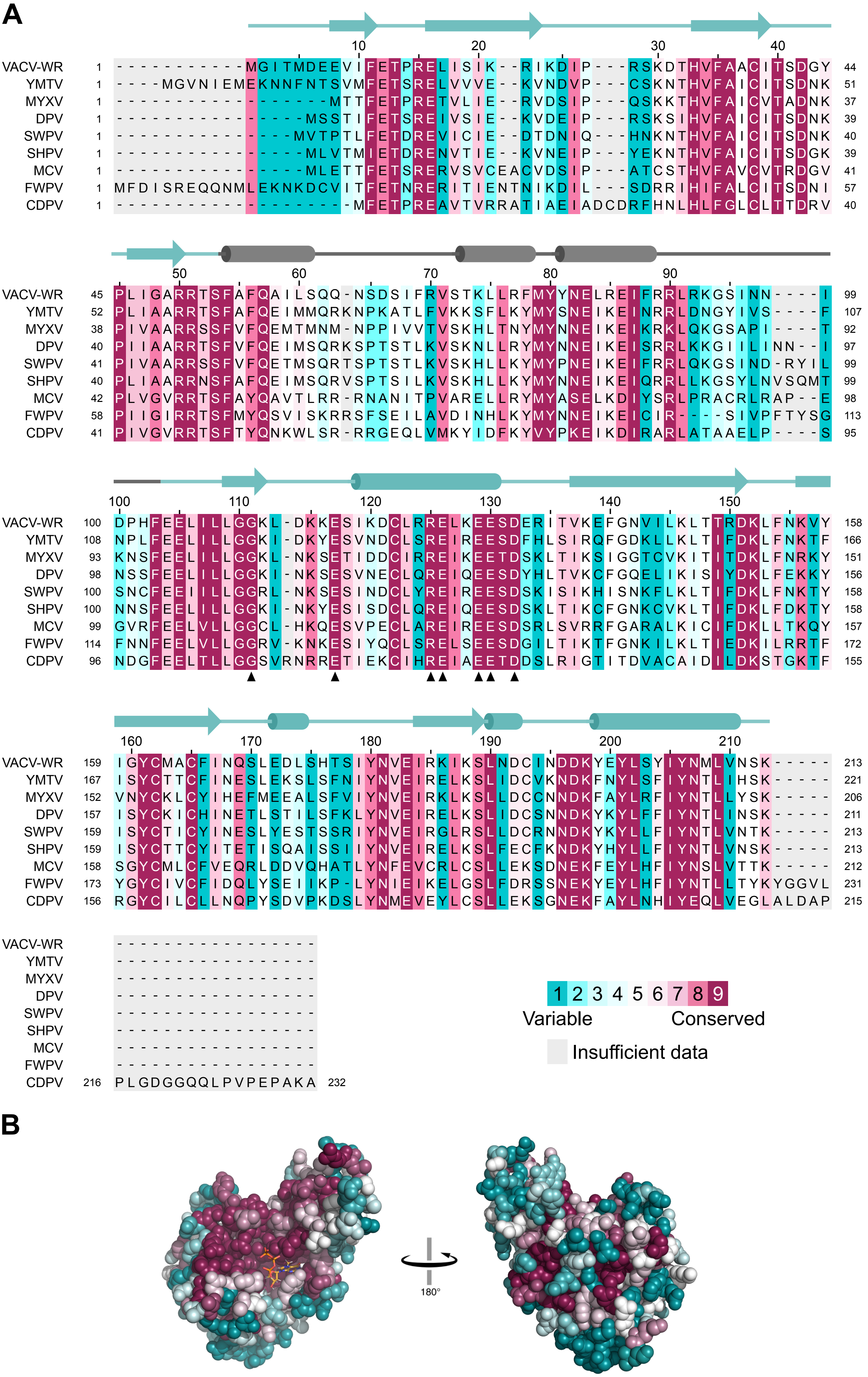

A
10 20 30 40
VACV-WR 1 -----------MGITMDEEVIFETPRELISIK--RIKDIP---RSKDTHVFAACITSDGY 44
YMTV 1 ----MGVNIEMEKNNFNTSVMFETSRELVVVE--KVNDVP---CSKNTHVFAICITSDNK 51
MYXV 1 ------------------MTTFETPRETVLIE--RVDSIP---QSKKTHVFAICVTADNK 37
DPV 1 ----------------MSSTIFETSREIVSIE--KVDEIP---RSKSIHVFAICITSDNK 39
SWPV 1 ---------------MVTPTLFETDREVICIE--DTDNIQ---HNKNTHVFAICITSDNK 40
SHPV 1 ----------------MLVTMIETDRENVTIE--KVNEIP---YEKNTHVFAICITSDGK 39
MCV 1 ----------------MLETTFETSRERVSVCEACVDSIP---ATCSTHVFAVCVTRDGV 41
FWPV 1 MFDISREQQNMLEKNKDCVITFETNRERITIENTNIKDIL---SDRRIHIFALCITSDNI 57
CDPV 1 --------------------MFETPREAVTVRRATIAEIADCDRFHNLHLFGLCLTTDRV 40
50 60 70 80 90
VACV-WR 45 PLIGARRTSFAFQAILSQQ-NSDSIFRVSTKLLRFMYYNELREIFRRLRKGSINN----I 99
YMTV 52 PLIAARRTSFAFQEIMMQRKNPKATLFVKKSFLKYMYSNEIKEINRRLDNGYIVS----F 107
MYXV 38 PIVAARRSSFVFQEMTMNM-NPPIVVTVSKHLTNYMYNNEIKEIKRKLQKGSAPI----T 92
DPV 40 PIIAARRTSFVFQEIMSQRKSPTSTLKVSKNLLRYMYNNEIKEISRRLKKGIILINN--I 97
SWPV 41 PIVAARRSSFVFQETMSQRTSPTSTLKVSKHLLKYMYPNEIKEIFRRLQKGSIND-RYIL 99
SHPV 40 PLIAARRNSFAFQEIMSQRVSPTSILKVSKHLLKYMYNNEIKEIQRRLLKGSYLNVSQMT 99
MCV 42 PLVGVRRTSFAYQAVTLRR-RNANITPVARELLRYMYASELKDIYSRLPRACRLRAP--E 98
FWPV 58 PIIGIRRTSFMYQSVISKRRSFSEILAVDINHLKYMYNNEIKEICIR---SIVPFTYSG 113
CDPV 41 PIVGVRRTSFTYQNKWLSR-RRGEQLVMKYIDFKYVYPKEIKDIRARLATAAELP----S 95
100 110 120 130 140 150
VACV-WR 100 DPHFEELILLGGKL-DKKESIKDCLRRELKEESDERITVKEFGNVILKLTTRDKLFNKVY 158
YMTV 108 NPLFEELILLGGKI-DKYESVNDCLSREIREESDFHLSIRQFGDKLLKLTIFDKLFNKTF 166
MYXV 93 KNSFEELILLGGKL-NKSETIDDCIRREIKEETDSKLTIKSIGGTCVKITITDKLFNRKY 151
DPV 98 NSSFEELILLGGKI-NKSESVNECLQREIQEESDYHLTVKCFGQELIKISIYDKLFEKKY 156
SWPV 100 SNCFEEIILLGGRL-NKSESINDCLYREIREESDSKISIKHISNKFLKLTIFDKLFNKTY 158
SHPV 100 NNSFEELILLGGKI-NKYESISDCLQREIQEESDSKLTIKCFGNKCVKLTIFDKLFDKTY 158
MCV 99 GVRFEELVLLGGCL-HKQESVPECLAREIREESDSRLSVRRFGARALKICIFDKLLTKQY 157
FWPV 114 FNNFEELVLLGGRV-KNKESIYQCLSRELSEESDGILTIKTFGNKILKLTIEDKILRRTF 172
CDPV 96 NDGFEELTLLGGSVRNRRETIEKCIHREIAEETDDSLRIGTITDVACAIDILDKSTGKTF 155
160 170 180 190 200 210
VACV-WR 159 IGYCMACFINQSLEDLSHTSIYNVEIRKIKSLNDCINDDKYEYLSYIYNMLVNSK----- 213
YMTV 167 ISYCTTCFINESLEKSLSFNIYNVEIRELKSLIDCVKNDKFNYLSFIYNTLIHSK----- 221
MYXV 152 VNYCKLCYIHEFMEEALSFVIYNVEIRKLKSLLDCCNNDKFAYLRFIYNTLLYSK----- 206
DPV 157 ISYCKICHINETLSTILSFKLYNVEIRELKSLIDCSNNDKYKYLFFIYNTLINSK----- 211
SWPV 159 ISYCTICYINESLYESTSSRIYNVEIRGLRSLLDCRNNDKYKYLLFIYNTLVNTK----- 213
SHPV 159 ISYCTTCYITETISQAISSIIYNVEIRELKSLFECFKNDKYHYLLFIYNTLVNSK----- 213
MCV 158 SGYCMLCFVEQRLDDVQHATLYNFEVCRLCSLLEKSDNEKFEYLHFIYNSLVTTK----- 212
FWPV 173 YGYCIVCFIDQLYSEIIKP-LYNIEIKELGSLFDRSSNEKYEYLHFIYNTLLTYKYGGVL 231
CDPV 156 RGYCILCLLNQPYSDVPKDSLYNMEVEYLCSLLEKSGNEKFAYLNHIYEQLVEGLALDAP 215
VACV-WR -----------------
YMTV -----------------
MYXV -----------------
DPV -----------------
SWPV -----------------
SHPV -----------------
MCV -----------------
FWPV -----------------
CDPV 216 PLGDGGQQLPVPEPAKA 232
1 2 3 4 5 6 7 8 9
Variable
Conserved
Insufficient data
B
180°

#### Figure S2

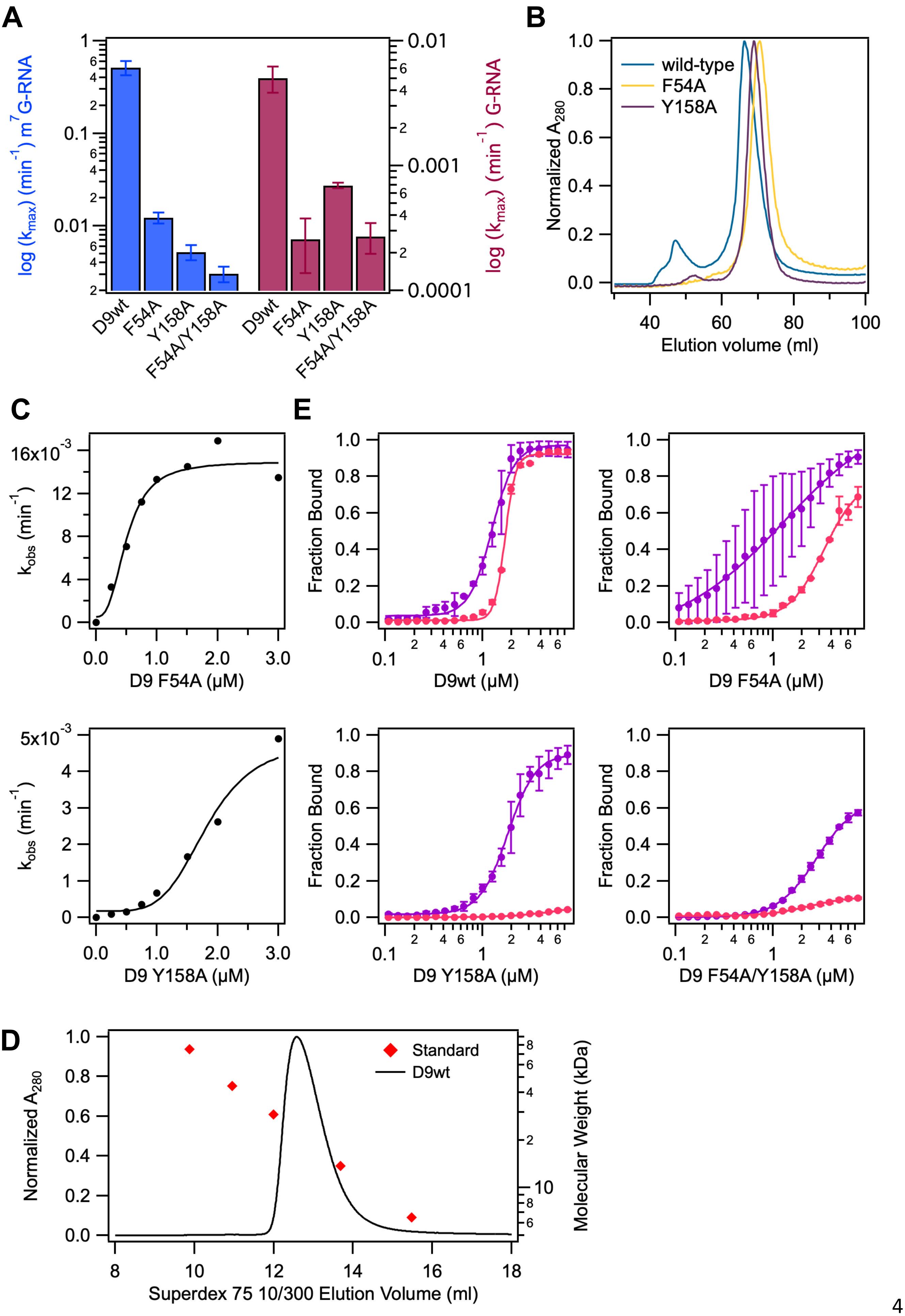
A
log (k_max) (min^-1) m^7G-RNA
log (k_max) (min^-1) G-RNA
D9wt
F54A
Y158A
F54A/Y158A
B
Normalized A280
wild-type
F54A
Y158A
Elution volume (ml)
C
k_obs (min^-1)
D9 F54A (μM)
D9 Y158A (μM)
E
Fraction Bound
D9wt (μM)
D9 F54A (μM)
D9 Y158A (μM)
D9 F54A/Y158A (μM)
D
Normalized A280
Standard
D9wt
Molecular Weight (kDa)
Superdex 75 10/300 Elution Volume (ml)

### Figure S3

**A**

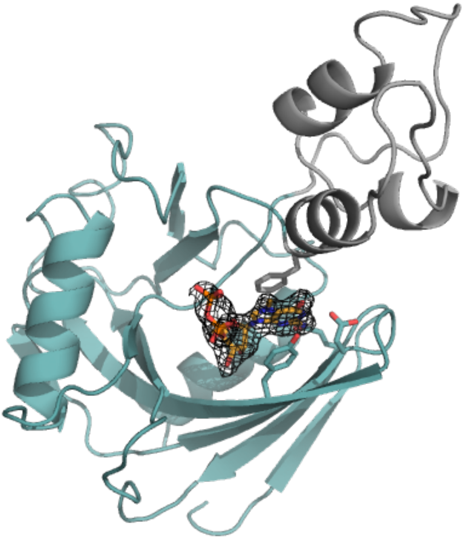

**B**

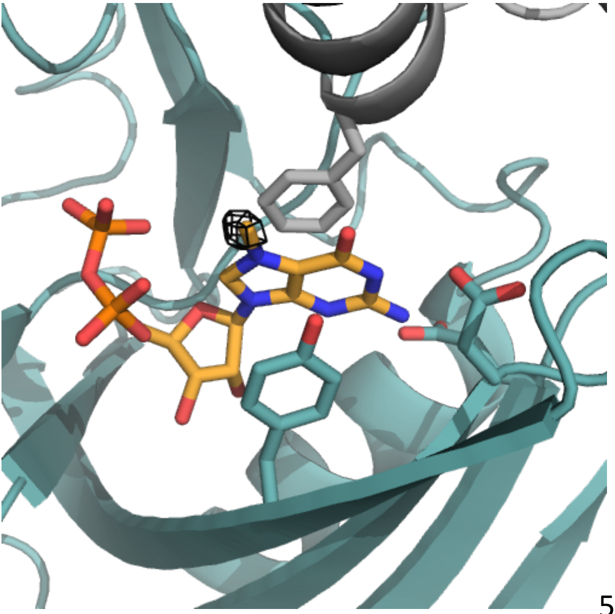

### Figure S4

## A

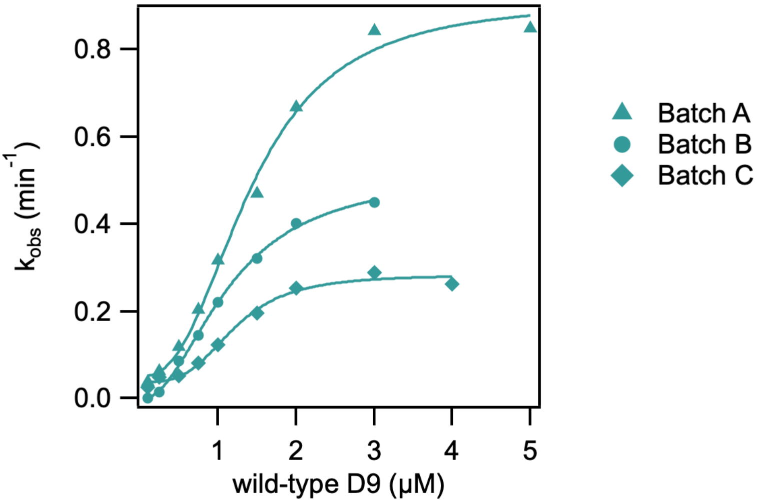


## B

| Preparation | $K_m$ (μM) | $k_{max}$ ($min^{-1}$) | Hill coefficient |
|---|---|---|---|
| Batch A | 1.42 ± 0.092 | 0.912 ± 0.044 | 2.5 |
| Batch B | 1.16 ± 0.061 | 0.517 ± 0.023 | 2.0 |
| Batch C | 1.21 ± 0.080 | 0.284 ± 0.013 | 3.4 |

**Figure S5**

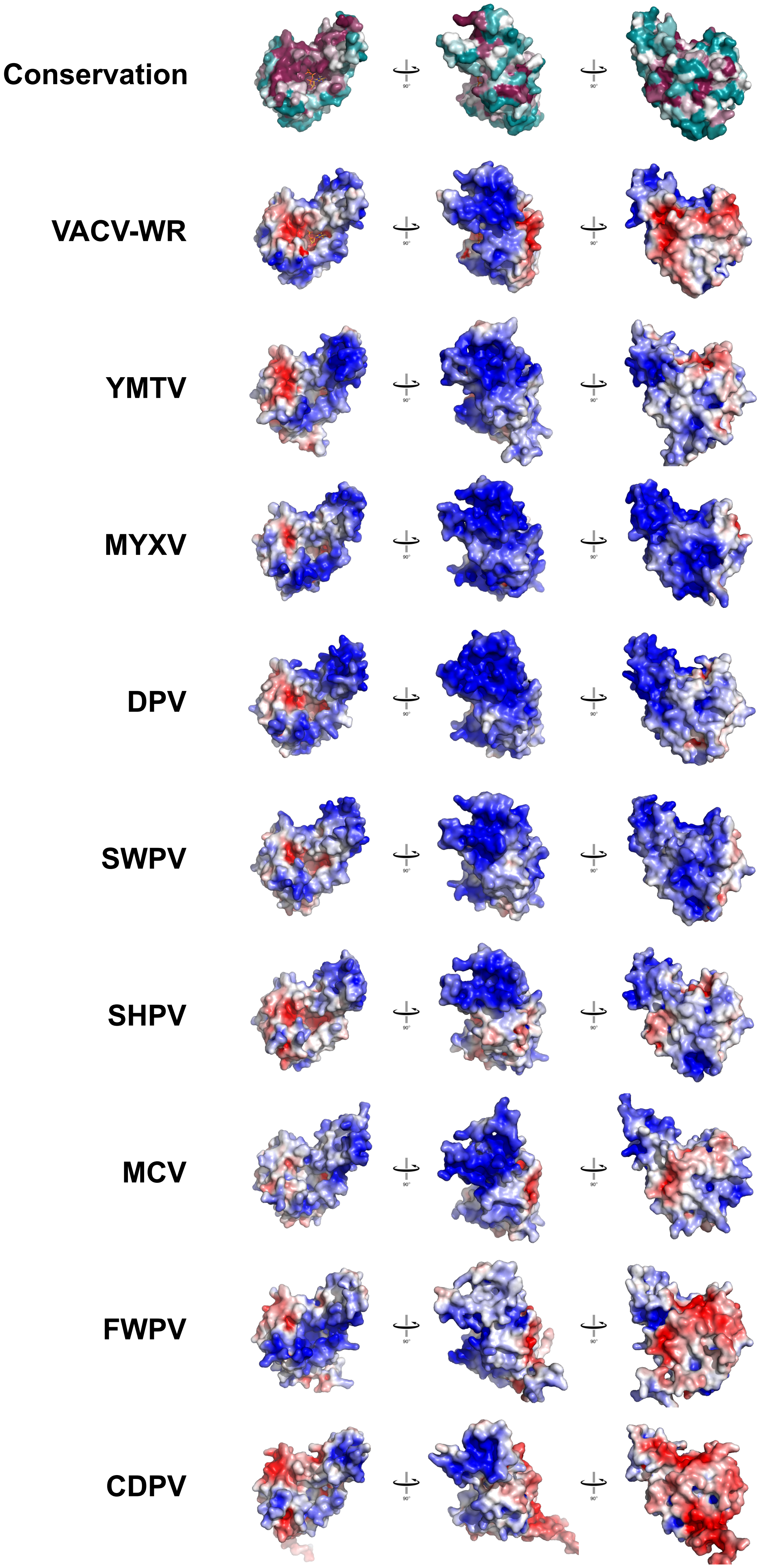

Tabel S1. Substrate binding affinities measured by filter binding assay.

| D9 construct | RNA | $K_d$ (μM) | Hill Coefficient |
|---|---|---|---|
| wt | $m^7G$-$GN_{28}$ | 1.25 | 3.7 |
| | G-$GN_{28}$ | 1.56 | 4.2 |
| | $m^7G$-$AN_{28}$ | 1.41 | 6.4 |
| | $m^7G$-$m^6AN_{28}$ | 1.27 | 3.7 |
| | p-$GN_{28}$ | 1.71 | 8.08 |
| | 5' overhang dsRNA | 0.87 | 6.0 |
| | Blunt end dsRNA | 1.32 | 9.0 |
| | 3' overhang dsRNA | 1.29 | 10.3 |
| F54A | $m^7G$-$GN_{28}$ | 1.28 | 0.78 |
| | p-$GN_{28}$ | 3.37 | 2.38 |
| Y158A | $m^7G$-$GN_{28}$ | 1.82 | 3.04 |
| | p-$GN_{28}$ | n.b.[a] | - |
| F54A/Y158A | $m^7G$-$GN_{28}$ | 2.82 | 2.17 |
| | p-$GN_{28}$ | n.b.[a] | - |

[a]Binding affinities could not be determined for lack of sufficient binding at highest protein concentration.

Table S2. RNA sequences used for activity and binding assays.

| Name | RNA sequence |
|---|---|
| $GN_{28}$ | GGGCAGAUUACAGCACGACAUAACACUAA (29)[a] |
| $AN_{28}$ | AGGCAGAUUACAGCACGACAUAACACUAA (29) |
| $m^6AN_{28}$ | $m^6$AGGCAGAUUACAGCACGACAUAACACUAA (29) |
| 5' overhang AS RNA | **GGG**UUAGUGUUAUGUCGUGCUG (22) |
| Blunt capped end AS RNA | **GGG**UUAGUGUUAUGUCGUGCUGUAAUCUGCCC (32) |
| 3' overhang AS RNA | **GGG**UUAGUGUUAUGUCGUGCUGUAAUCUGCCC**AAUAACUACC** (42) |

[a]Values in parentheses denote RNA length. Nucleotides in bold do not anneal to complementary RNA ($GN_{28}$) in dsRNA constructs.
